# Rapid Progesterone signaling induces meiosis through a dual coreceptor trafficking trap

**DOI:** 10.64898/2026.10.27.762173

**Authors:** Lama Assaf, Nancy Nader, Mustafa Shammout, Hana Mohamad, Sama Al-Saffar, Khaled Machaca

## Abstract

Agonist-dependent signaling through plasma membrane (PM) receptors takes place through diverse modes, including post-translational modifications, second messengers, formation of signaling complexes, or ionic fluxes among others. Classical signaling downstream of steroid hormones engages nuclear/cytoplasmic receptors to induce transcription. Steroids further signal through a rapid mode within seconds to minutes independently of transcription. Despite the now well-recognized physiological roles of rapid progesterone (P4) signaling, the mechanistic steps remain poorly defined. Here we elucidate the early P4 rapid signaling steps that mediate the release of the oocyte meiotic arrest in the frog. P4 signals through two coreceptors mPRβ, which localizes at the PM; and ABHD2, which localizes intracellularly. Binding of P4 increases the interaction affinity between the two coreceptors leading to trapping of ABHD2 at the PM and activating signaling. Although P4 is not required for mPRβ-ABHD2 interaction at artificially high receptor concentration, it is essential under physiological conditions. We further show that ABHD2 is a dual pass transmembrane protein that binds P4 leading to intracellular PLA2 activity. Signaling through several receptor classes involves endocytosis, but the mPRβ-ABHD2 signaling modality is novel as it employs a PM trafficking trap to form the coreceptor complex and induce meiosis. Hence, our findings define a novel signaling modality for rapid steroid signaling.

**Significance Statement:** Define the early rapid progesterone signaling steps involving a trafficking trap at the PM due to increased coreceptor interaction affinity to trigger lipase activation to induce meiosis resumption.

## Introduction

Plasma membrane (PM) receptors respond to agonists by inducing signaling cascades that are rarely linear and often branch in multiple directions to mediate their biological action. Signal branching allows for better integration and thus adaptation of the cell to its environment. As cells are exposed to various agonists and internal modulators, it is useful to think of agonist action as tipping the balance in one direction or the other rather than an on-off switch.

One such signaling cascade is the release of the prophase I meiotic arrest in *Xenopus laevis* oocytes, which is mediated by progesterone (P4) through a rapid non-genomic signaling pathway. This is the first step in preparing the egg for fertilization and embryonic development. Fully grown vertebrate oocytes are arrested at prophase of meiosis I but cannot be fertilized until they undergo oocyte maturation ^1^. Maturation encompasses both nuclear and cytoplasmic maturation. Nuclear maturation involves meiosis progression with the extrusion of the first polar body and arrest at metaphase of meiosis II. Cytoplasmic maturation leads to remodeling of the oocyte cellular signaling pathways and organelles in preparation for fertilization ^2–5^.

Classical P4 signaling is mediated through nuclear receptors (nPR) that induce transcription of gene subsets to encode specific biological functions ^6,7^. P4 in addition signals at a more rapid time scale without a direct reliance on transcription -although in some cases it may result in transcriptional changes-through a mechanism referred to as rapid or non-genomic signaling. In contrast to the nPRs that act as transcription factors, rapid P4 signaling relies on membrane progesterone receptors (mPR), which are high affinity PM serpentine receptors that belong to the progestin and AdipoQ receptors family (PAQR). There are 11 PAQR receptors in vertebrates with five that have been shown to respond to P4: PAQR 5, 6, 7, 8, and 9 (mPR γ, δ, α, β, and ɛ) ^8^.

Since the cloning of the first mPR from fish ovaries over two decades ago ^9^, evidence is accumulating for important roles for the different mPR isoforms in the nervous system, reproductive tissues, cardiovascular system, immunity, cancer, and metabolism ^10–14^. This fits with the broad tissue distribution of mPR isoforms; however, the specific roles of distinct mPRs are not well defined apart from mPRα, which is the most heavily investigated ^12,15^.

In addition to mPRs, other putative receptors have been implicated in rapid P4 signaling, although with mixed supporting evidence. Those can be classified broadly in three families: 1) membrane-associated P4 receptors (MAPRs), represented by four members with widely diverse physiological functions, including PGRMC1 and 2 (Progesterone Receptor Membrane Component 1 and 2), which are small heme-binding proteins that interact with cytochrome P450 ^16,17^. 2) Neurosteroid receptors, such as GABA_A_ and NMDA receptors, which respond to P4 and its metabolites, including allopregnanolone, in the brain ^17^. 3) ABHD2 (Abhydrolase Domain Containing 2), which is a proposed P4 receptor in sperm that acts as a monoacylglycerol lipase (MAGL) to activate the chemosensory Ca^2+^ channel CatSper leading to hypermotility ^18^. ABHD2 mouse knockout lines reveal roles in vascular smooth muscle and lung function with implications on atherosclerosis and emphysema ^19,20^. ABHD2 is also highly expressed in the ovary, and its ablation leads to dysregulation of the estrus cycle ^21^.

Studying rapid P4 signaling is complicated by the multitude of potential receptors, their tissue expression profile, affinity, and the metabolism of P4 itself, which can produce additional potential agonists. Oocyte maturation in the anuran *Xenopus laevis* has been a historically attractive system to study rapid P4 signaling because the oocyte is transcriptionally silent thus minimizing the likelihood of a role for nPRs; and because a single mPR isoform, mPRβ (PAQR8), is required for meiosis resumption ^22,23^.

mPRβ has the opposite topology to that of GPCRs with an intracellular N-terminus and an extracellular C-terminus and is internalized into an endosomal compartment in response to P4 in the oocyte ^23^. Furthermore, mPRβ signaling requires the adaptor protein APPL1 and the kinase Akt2 to initiate the kinase cascade that leads to MPF activation and the progression of meiosis ^23^.

Recently, we showed that in addition to mPRβ, ABHD2 is also essential for P4-dependent oocyte maturation ^24^ (Fig. 1A). Interestingly, mPRβ and ABHD2 act as obligate co-receptors, with both required for P4 action ^24^. Overexpressing either receptor does not rescue the loss of the other. Furthermore, in the oocyte ABHD2 exhibits PLA2 activity, resulting in the generation of prostaglandins, lysophosphatidic acid (LPA), and sphingosine 1 phosphate to support oocyte maturation ^24^. Importantly, the ABHD2 PLA2 activity requires both the catalytic residues in its lipase domain and interaction with mPRβ as a co-receptor ^24^. It is however not clear how P4 induces mPRβ and ABHD2-dependent signaling. Furthermore, the requirement for a co-receptor during mPR signaling is novel, so understanding the signaling details is important.

**Figure 1.**
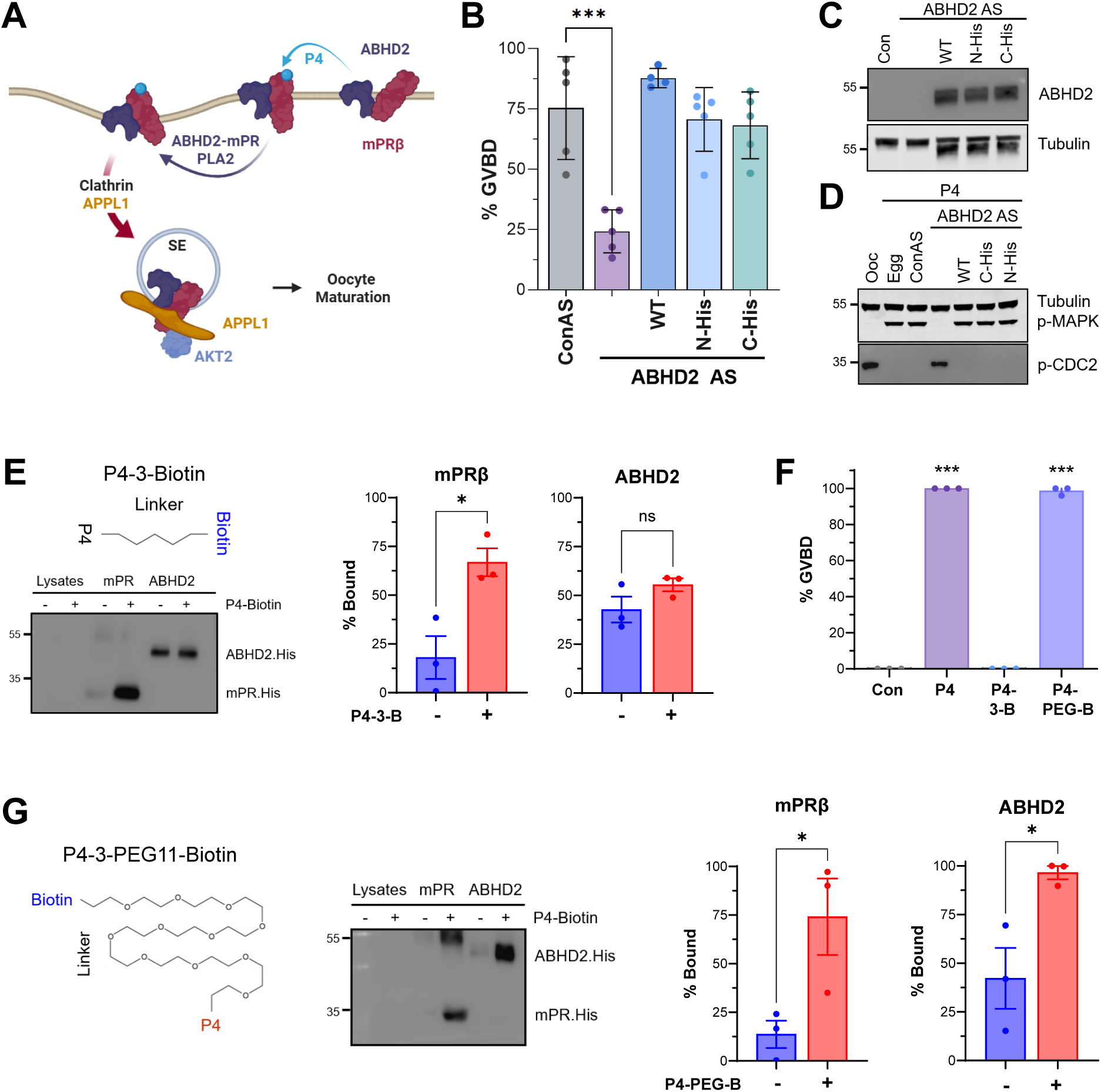
ABHD2 binds P4. **(A)** Current model of rapid P4 signaling in *Xenopus* oocytes. **(B)** Germinal vesicle breakdown (GVBD) control antisense (Con AS)-, or ABHD2 antisense (ABHD2 AS)-injected oocytes expressing untagged ABHD2 (WT), N-terminally His-tagged ABHD2 (His-ABHD2), or C-terminally His-tagged ABHD2 (ABHD2-His), following P4 stimulation. GVBD was normalized to naïve oocytes from the corresponding experiment. Data are presented as Mean ± SEM, n = 4-5, one-way ANOVA Dunnett correction comparing each group with Con AS (*** p < 0.001; not significant comparisons are not shown). **(C-D)** Representative Western blots showing the expression of the ABHD2 constructs and P4-induced MAPK phosphorylation (C) and Cdc2 dephosphorylation (D). Tubulin serves as a loading control. Blots are representative of three independent experiments. **(E)** Structure of P4-3-biotin containing a short linker between P4 and biotin. Binding of P4-3-B to mPRβ or ABHD2 expressed in microsomes. Microsomal lysates were incubated with P4-3-B (10^−5^ M) or vehicle, followed by streptavidin pulldown and Western blotting using an anti-His antibody. Representative Western blot and quantification of bound proteins are shown. The percent bound was calculated from band intensities as IP/(IP + FT) × 100, where IP and FT represent the streptavidin-bound and flow-through fractions, respectively. Data are presented as Mean ± SEM, n = 3, unpaired two-tailed t-test (* p < 0.05; ns, not significant). **(F)** Percent GVBD in oocytes treated with vehicle control or 10^−5^ M P4, P4-3-Biotin (P4-3-B), or P4-3-PEG11-biotin (P4-PEG-B). Data are presented as Mean ± SEM, n = 3, ANOVA with Tukey correction (*** p < 0.001; not significant not shown). **(G)** Structure of P4-PEG-B with a longer PEG11 linker. Binding of P4-PEG-B to mPRβ or ABHD2 expressed microsomes. Microsomal lysates were incubated with P4-PEG-B (10^−5^ M) or vehicle, followed by streptavidin pulldown and Western blotting using an anti-His antibody. Representative Western blot and quantification of bound proteins quantified as described in (E) are shown. Data are presented as Mean ± SEM, n = 3, two-tailed t-test (* p < 0.05).

Here we show that at steady state ABHD2 is enriched intracellularly whereas mPRβ is mostly at the PM. ABHD2 continuously samples the PM through recycling. P4 increases the affinity of interaction between mPRβ and ABHD2 resulting in the trapping of the coreceptor complex at the PM and triggering its PLA2 activity. Increasing the levels of the two receptors within the same vesicular compartment both in vivo and in vitro is sufficient to bypass the requirement for P4. We further show that ABHD2 is a two-pass transmembrane protein that binds P4 leading to intracellular PLA2 activity. Although several receptor classes, including GCPRs and TGFβ receptors, involve endocytosis in their signaling, the activation mechanism of the mPRβ-ABHD2 coreceptor complex represents a novel mode of agonist receptor signaling where a trafficking trap recruits the coreceptor complex to the PM to trigger signaling.

## Results

### ABHD2 binds P4

To better understand the mechanisms regulating mPRβ-ABHD2 signaling, we wanted to track the subcellular location of ABHD2. We had previously tagged ABHD2 at the N- or C-terminus with mCherry but neither construct rescued oocyte maturation following ABHD2 knockdown ^24^. As this could be due to steric hindrance by the large mCherry tag, we placed mCherry before the transmembrane domain and separated it from ABHD2 using either rigid or flexible linkers (Fig. S1A). The constructs were expressed in both cases (Fig. S1B), but they did not rescue oocyte maturation following ABHD2 knockdown (Fig. S1C and S1D). This argues that ABHD2 has little tolerance for a large tag at either terminus, suggesting that precise ABHD2 topology is important.

We therefore tested the smaller His-tag. ABHD2 tagged with His at either the N- or C-terminus is functional and rescues oocyte maturation following ABHD2 knockdown (Fig. 1B). Both the His-ABHD2 and the ABHD2-His clones expressed to similar levels to the wild type untagged ABHD2 (WT) (Fig. 1C). Furthermore, expression of the three constructs led to phosphorylation of MAPK and dephosphorylation of Cdc2 in response to P4 (Fig. 1D), showing proper activation of the kinase cascade driving oocyte maturation ^25^.

Mechanistically the mPRβ-ABHD2 coreceptor complex could be assembled and activated by P4 binding to either mPR alone or to both mPR and ABHD2. Both mPRα and human ABHD2 have been reported to bind P4 ^18,26^. However, a recent report using recombinant human ABHD2 argues that it does not bind P4 ^27^. To test the capacity of *Xenopus* ABHD2 to bind P4, we expressed ABHD2-His in the tobacco lysate system (ALiCE) as it produces high protein yields in microsomes for biochemical analyses ^24^. We incubated the microsomes with P4-3-Biotin (Fig. S2A) and purified the complex on streptavidin beads, using His-tagged mPR as a positive control (Fig. 1E and S2C). mPR bound to the beads only when the microsomes were pre-incubated with P4-3-Biotin (Fig. 1E and S2C) and was competed off the beads with free P4 (Fig. S2D). This shows that mPRβ binds to P4-3-Biotin. In contrast, ABHD2 did not bind to P4-3-Biotin as we did not observe significantly increased binding to the streptavidin beads when the microsome were incubated with P4-3-Biotin (Fig. 1E). Although in this case ABHD2 exhibited higher non-specific binding to the beads (Fig. 1E and S2C) that could not be competed with free P4 (Fig. S2E). These results show that ABHD2 is unable to bind P4-3-Biotin.

However, as the P4-3-Biotin has a short linker (Fig. 1E and S2A), we wanted to confirm that it is functional biologically to rule out the possibility of the biotin moiety interfering with P4’s ability to bind to the receptor(s) in vivo. Interestingly, P4-3-Biotin was unable to induce oocyte maturation at similar concentrations to free P4 (Fig. 1F), showing that it’s not active biologically. To test whether this is due to the short linker, we assessed the activity of a P4-Biotin with a longer linker P4-3-PEG11-Biotin (Fig. 1F and S2B). P4-3-PEG11-Biotin induced maturation to similar levels as free P4 (Fig. 1F), showing that the short linker in P4-3-Biotin interferes with the ability of P4 to induce oocyte maturation. As P4-3-PEG11-Biotin is active in vivo, we tested whether it binds to ABHD2 expressed in microsomes. As shown in Figure 1G, P4-3-PEG11-Biotin binds to both mPRβ and ABHD2 (Fig. 1G and S2F). Therefore, the ability of the biotin-labelled P4 to induce maturation correlates with their ability to binds ABHD2. This not only shows that P4 binds ABHD2 but that this binding is required to induce oocyte maturation.

### ABHD2 topology

We previously showed that the oocyte produces various lipid metabolites downstream of PLA2 in response to P4 ^24^, arguing that they are generated intracellularly. We were thus interested in visualizing the mPRβ-ABHD2 coreceptor complex PLA2 activity. We injected a fluorescently labelled PLA2 substrate (PED6) to report PLA2 activity live in the oocyte. We observed a time-dependent increase in PED6 fluorescence only after P4 addition (Fig. 2A). Importantly this increase required ABHD2 as it was lost in oocytes when ABHD2 was knocked down (Fig. 2A). These results support our lipidomic studies ^24^, and suggest that the ABHD2-dependent PLA2 activity in the oocyte produces cytosolic metabolites by acting on the inner leaflet of the PM.

**Figure 2.**
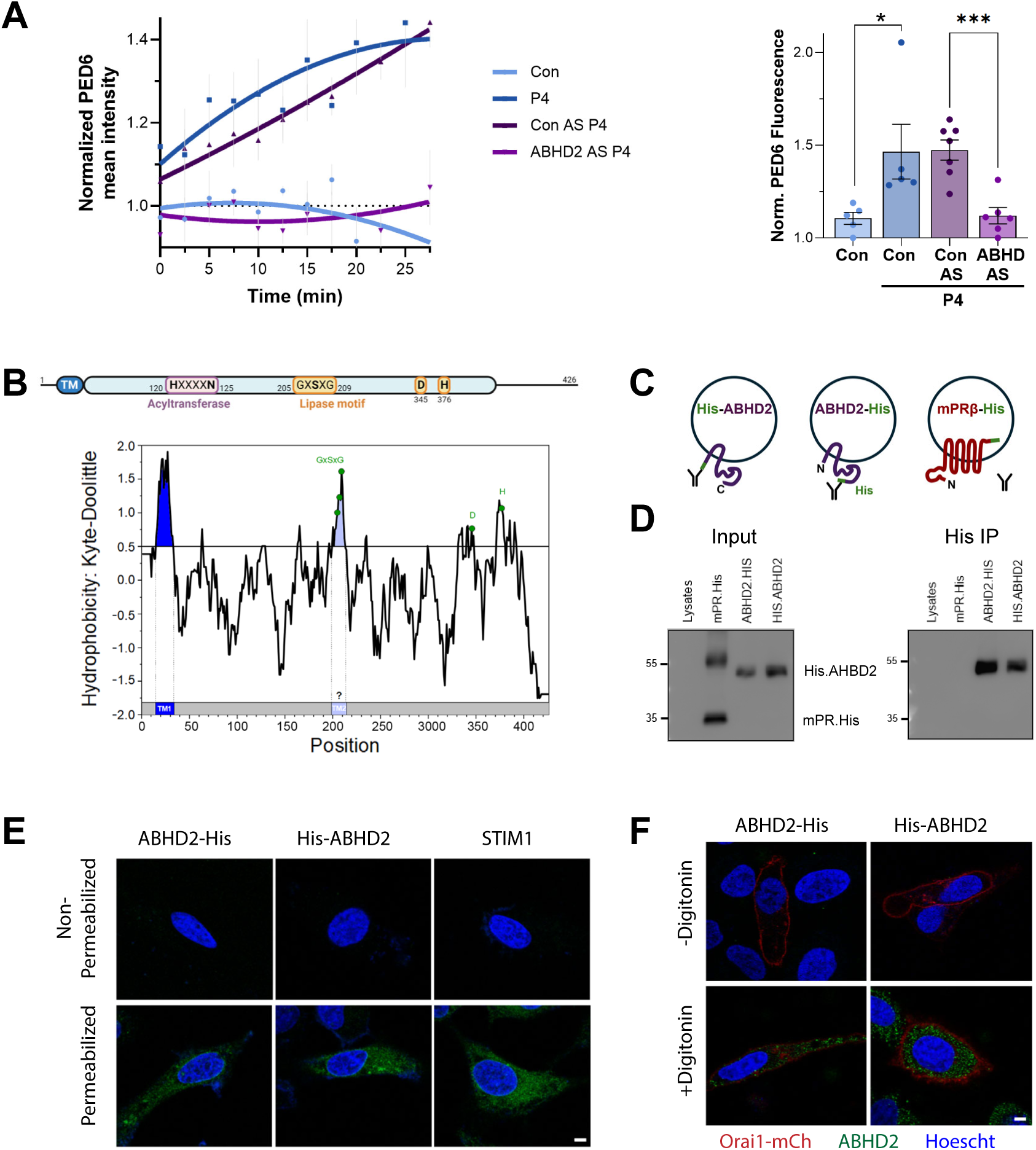
ABHD2 Membrane Topology. **(A)** Live-cell measurement of PLA, activity in oocytes microinjected with the fluorescent PLA, substrate PED6 (10 µM). Oocytes were injected with control antisense (Con AS) or ABHD2 antisense (ABHD2 AS) followed by P4 (10^−5^ M) or ethanol treatment. PED6 fluorescence intensity in the oocyte cortex. Fluorescence values were normalized to the pretreatment baseline intensity of individual oocyte. Fluorescence time courses were fitted using second-order polynomial regression (left). The bar graph (right) represents normalized fluorescence measured at 30 min after treatment for each oocyte. Data are presented as Mean ± SEM, n = 5-7 oocytes from three donor females, unpaired two-tailed t-test (*p < 0.05; ***p < 0.001). **(B)** Schematic representation of *Xenopus* ABHD2 highlighting the predicted transmembrane domains (TM), acyltransferase motif (HXXXXN), and the conserved lipase motif (GxSxG, D345, and H376). Kyte–Doolittle hydropathy plot, generated using a nine-residue window, identifies two putative transmembrane regions with the second overlapping the conserved GxSxG lipase motif. **(C)** Schematic representation of the predicted membrane topology and His-tag accessibility of N-(His-ABHD2) or C-terminally (ABHD2-His) His-tagged ABHD2 and C-terminally His-tagged mPRβ (mPRβ-His) in microsomes. **(D)** Microsomes were immunoprecipitated with an anti-His antibody followed by Western blotting. Representative input and immunoprecipitated blot is shown. Representative of three independent experiments. **(E)** Immunofluorescence of His-ABHD2 or ABHD2-His expressed in HeLa cells. Cells were immunostained using an anti-His antibody under non-permeabilized conditions or following permeabilization with 0.3% Triton X-100. STIM1 immunostaining served as a positive control for membrane permeabilization. Images are representative of three independent experiments. Scale bar 5 µm. **(F)** HeLa cells co-expressing His-ABHD2 or ABHD2-His with Orai1-mCherry as a PM marker were immunostained under non-permeabilized conditions or following selective plasma membrane permeabilization with 0.01% digitonin. Orai1-mCherry is shown in red, ABHD2 (anti-His) in green, and nuclei counterstained with Hoechst in blue. Images are representative of four independent experiments. Scale bar 5 µm.

The predicted topology of ABHD2 is that of a typical type II transmembrane protein with a cytoplasmic N-terminus and a large extracellular portion that contains the catalytic domain (Fig. 2B, top), which would imply that it acts on outer leaflet phospholipids ^18,28^. A hydropathy plot of *Xenopus* ABHD2, which is highly conserved relative to human ABHD2 (Fig. S3A), with a window of 9 residues reveals a potential shorter second transmembrane domain (TM) of 12 residues as compared to the first more robustly predicted TM of 18 residues (Fig. 2B, bottom). This topology is intriguing as the predicted second TM encompasses part of the consensus lipase domain of ABHD2 (205-GxSXG-209, D345, H376) (Fig. 2B), which would place the catalytic domain within the core of the bilayer with access to the fatty acid tails.

To elucidate the topology of ABHD2, we expressed the N- or C-terminally His-tagged proteins in microsomes. If ABHD2 contains two TM domains, we expect that both its N- and C-termini would be cytosolic and thus accessible to antibody pulldown (Fig. 2C). In contrast, if ABHD2 has a single TM domain then only the N-terminus would be accessible as the C-terminus would be extracellular, that is shielded within the microsome lumen. As shown in Figure 2D, we were able to pulldown both His-ABHD2 and ABHD2-His from microsomes showing that both termini are cytosolic. This supports a two transmembrane topology for ABHD2 with both termini in the cytosol. As a control for the integrity of the microsomes, we performed a similar experiment with a C-terminally His-tagged mPRβ, which we have previously shown to be extracellular ^24^. The C-terminal His-tagged mPRβ was not accessible to the antibody pulldown as would be expected from its luminal (extracellular) localization (Fig. 2C and 2D).

To confirm the ABHD2 topology from the microsome experiment, we transfected HepG2 cells with His-ABHD2 or ABHD2-His and immunostained for the His-tag in non-permeabilized cells and cells permeabilized with Triton X-100 to allow antibody access to the cytosol. As a control for PM permeabilization, we stained for STIM1, an ER resident protein (Fig. 2E). We detected the His-tag only in permeabilized cells whether His-ABHD2 or ABHD2-His were expressed (Fig. 2E). This supports the conclusion that both ABHD2 termini are cytoplasmic. However, a caveat is that ABHD2 could theoretically localize intracellularly within vesicles and not at the PM. In that case even if both the N- and C-termini are extracellular they would only be detected after permeabilization. To rule out this possibility we selectively permeabilized the PM using digitonin. In this case we co-expressed the tagged ABHD2 constructs with the PM localized channel Orai1 to mark the cell membrane (Fig. 2F). Both N- and C-terminal ABHD2 His tags were detected only after digitonin treatment and not in non-permeabilized cells (Fig. 2F), indicating that both ABHD2 termini face the cytoplasm.

Together these results support an ABHD2 topology with two transmembrane domains and with cytosolic N- and C-termini. This topology places the ABHD2 lipase domain within the core of the PM and cytosolically (Fig. S3B), thus producing intracellular PLA2 metabolites.

### ABHD2 and mPRβ signal in cis topologically

An intriguing observation from co-expressing ABHD2 and mPRβ in same microsomes at high concentrations is that their PLA2 activity becomes P4-independent (Fig. 3A-C) ^24^. This argues that at high concentrations mass action supports ABHD2-mPRβ interaction and PLA2 activation independently of P4. This does not occur physiologically as presumably receptor concentrations are limiting. However, we can forcibly enrich mPRβ into endosomes experimentally in the oocyte, which is sufficient to induce maturation independently of P4 ^23,24,29,30^. As mPRβ continuously recycles at the PM, we can enrich it in endosomes by blocking exocytosis using a dominant negative SNAP25 mutant, while allowing endocytosis to proceed, which triggers maturation independently of P4 ^23,24,29,30^. These results support the conclusion that high enough concentrations of both receptors within the same topological compartment drive their interaction independently of P4.

**Figure 3.**
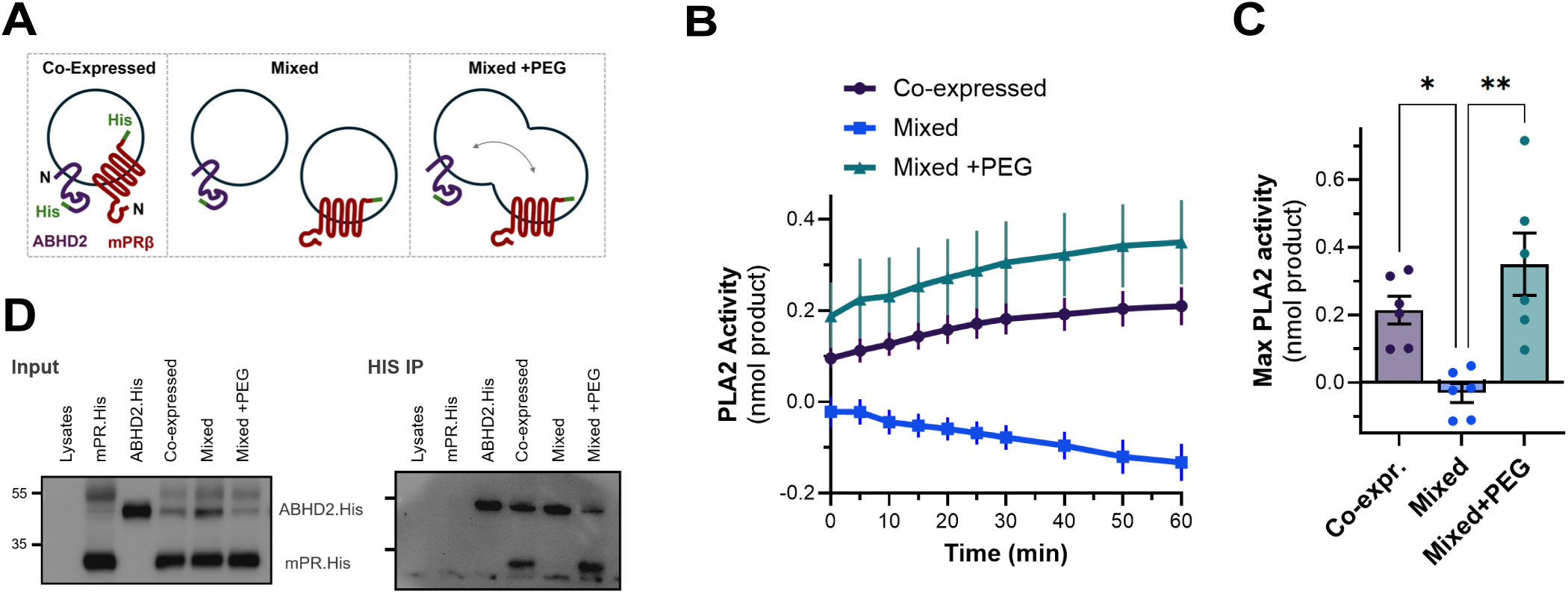
ABHD2 and mPRβ interact in cis. **(A)** Schematic representation of the experimental conditions used to assess the interaction of mPRβ and ABHD2. Both receptors were either co-expressed in the same microsomes (Co-expressed), expressed separately and subsequently mixed (Mixed), or expressed separately, mixed and treated with 10% polyethylene glycol (PEG) for 15 min at 37°C to induce microsome fusion (Mixed+PEG). **(B)** Time courses of PLA, activity in microsomes under the conditions illustrated in (A). PLA, activity was measured using a fluorometric assay with subtraction of the background from control lysates without DNA. Data are presented as Mean ± SEM, n = 6. **(C)** Quantification of the maximum PLA, product formed at 60 min. Data are presented as Mean ± SEM, n = 6, one-way ANOVA with Tukey correction (* p < 0.05; ** p < 0.01). **(D)** Representative Western blots of the input and anti-His immunoprecipitated fractions from microsomes expressing mPRβ-His and ABHD2-His as outlined in panel A. Blots are representative of four independent experiments.

To directly test this conclusion and further provide insights into whether ABHD2 and mPRβ function in cis or in trans topologically, we used the microsome expression system. Co-expression of ABHD2 and mPRβ in the same microsomes leads to P4-independent PLA2 activity (Fig. 3A-C, Co-expressed). However, when mPRβ and ABHD2 are expressed in separate microsomes and the microsome are then mixed, they do not support PLA2 activity (Fig. 3A-C, Mixed), arguing that the receptors must be within the same compartment topologically to function, that is they cannot interact in trans. To confirm this conclusion, we treated microsomes separately expressing mPRβ or ABHD2 with polyethylene glycol (PEG) as a crowding agent to induce microsome fusion to bring the two receptors together within the same compartment (Fig. 3A). PEG treatment endows the mixed microsomes with PLA2 activity (Fig. 3B-C, Mixed+PEG), supporting the conclusion that the two receptors must be within the same compartment to interact.

To support the functional PLA2 activity, we pulled down mPRβ-His or ABHD2-His from the three different microsome populations. As expected mPRβ-His was not accessible for pulldown as the tag is lumenal (Fig. 3D), whereas ABHD2-His is accessible and is pulled down from the microsomes under all conditions (Fig. 3D). Pulling down ABHD2-His co-precipitates mPRβ only in microsomes co-expressing both receptors and in PEG-treated mixed microsomes (Fig. 3A and 3D). mPR did not co-immunoprecipitate with ABHD2 in mixed microsomes in the absence of PEG (Fig. 3D), confirming that the two receptors are in separate microsome and do not interact. Collectively, these results show that both receptors must be within the same compartment to physically interact and mediate PLA2 activity.

### P4 increases the interaction affinity of mPRβ to ABHD2

The above results raise a question about the role of P4 in mediating the mPRβ-ABHD2 interaction. The simplest explanation is that P4 binding to the receptors increases their interaction affinity with each other to induce PLA2 activity at limiting receptors concentrations. To test this idea, we co-expressed both receptors in microsome and titrated ABHD2 with increasing mPR concentrations (Fig. 4A and 4B). We maintained ABHD2 DNA concentration in the expression system constant and gradually increased mPRβ DNA (2.5, 5, and 10 nM) (Fig. 4A, top). At the lower concentrations of mPRβ (2.5 and 5 nM) PLA2 activity was P4-dependent, whereas at the higher concentration (10 nM) it was not (Fig. 4A and 4B). Expression analysis confirms the levels of the two receptors in the microsomes and shows a shift in their ratio, as apparently increasing mPRβ DNA not only increases mPRβ protein expression but also lowers the amount of expressed ABHD2 presumably due to taxing the transcription/translation machinery in the expression system (Fig. 4C).

**Figure 4.**
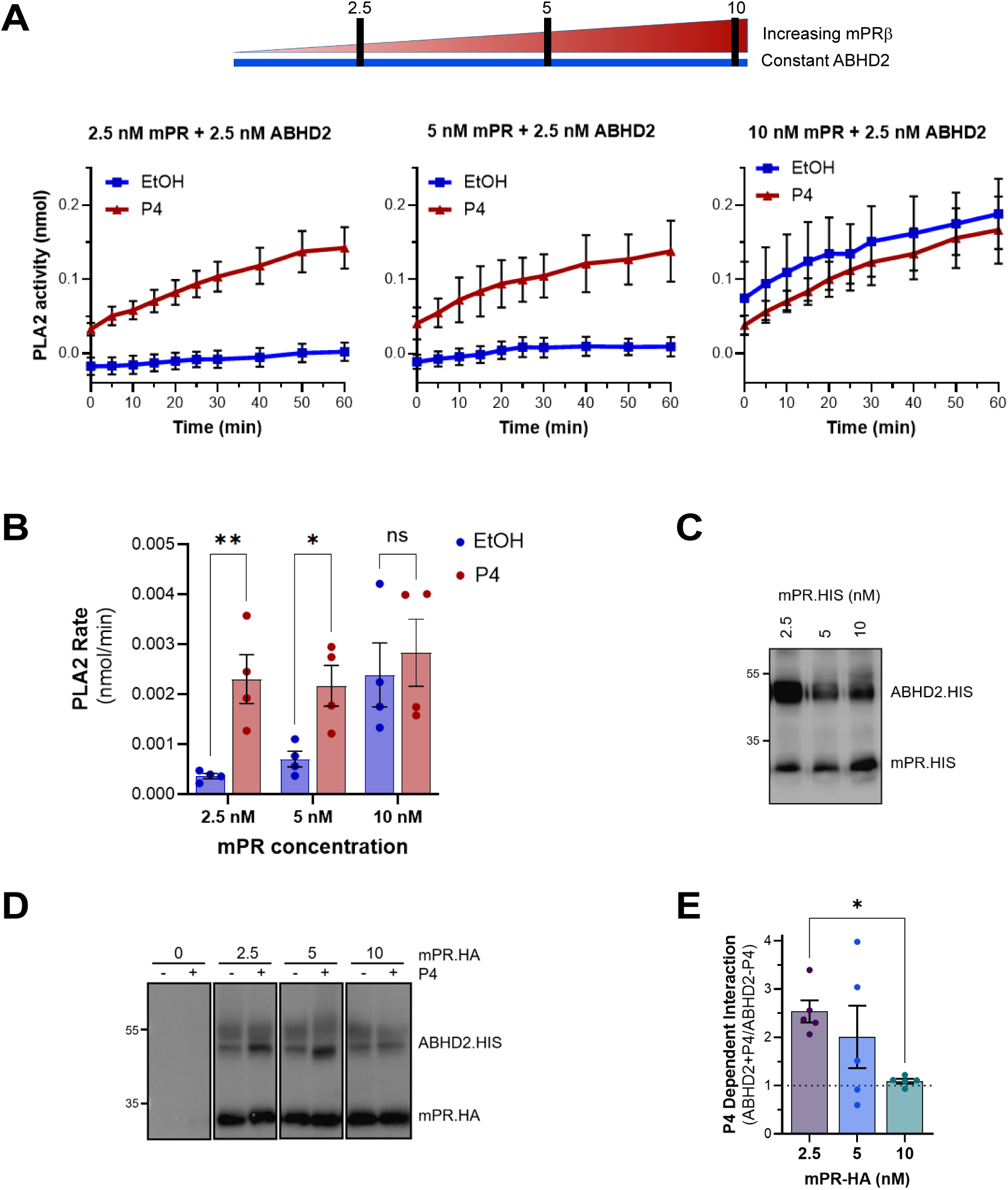
P4 enhances mPRβ–ABHD2 interaction affinity. **(A)** Schematic representation of increasing mPRβ DNA concentrations (2.5, 5, and 10 nM) while maintaining a constant ABHD2 DNA concentration (2.5 nM). PLA, activity was measured in microsomes co-expressing mPRβ and ABHD2 at the indicated concentrations following treatment with the ethanol control (EtOH) or P4 (10^−5^ M). Data are presented as Mean ± SEM, n=4. **(B)** PLA, rate quantified as the slope of the fluorescence time courses in (A). Data are presented as Mean ± SEM, n = 4, two-way ANOVA with Šídák’s correction (* p < 0.05; ** p < 0.01; ns, not significant). **(C)** Representative Western blot of mPRβ-His and ABHD2-His expression in microsomes at increasing mPRβ DNA concentrations (2.5, 5, and 10 nM), while maintaining ABHD2 DNA constant at 2.5 nM. **(D)** Representative anti-HA co-immunoprecipitation of ABHD2-His with HA-tagged mPRβ expressed at increasing DNA concentrations (0, 2.5, 5, and 10 nM) in microsomes treated with P4 (10^−5^ M) or ethanol. ABHD2-His was maintained at a constant DNA concentration of 2.5 nM. Blots are representative of five independent experiments. **(E)** Quantification of P4-dependent ABHD2–mPRβ association from the co-immunoprecipitation experiments shown in (D). Co-immunoprecipitated ABHD2-His band intensities were quantified and expressed as the ratio in the presence over the absence of P4 (+P4/−P4) at each mPRβ DNA concentration. Data are presented as Mean ± SEM, n = 5, one-way ANOVA with Tukey correction (* p < 0.05).

To confirm that the observed PLA2 activity reflects and depends on the levels of mPRβ-ABHD2 interaction, we pulled down HA-mPRβ from the microsomes. As the expression levels of mPRβ increased the P4-dependent interaction between ABHD2 and mPRβ decreased (Fig. 4D and 4E). At the lower mPRβ concentrations (2.5 and 5 nM) P4 enhanced the interaction between mPRβ and ABHD2 (Fig. 4D and S4A), whereas at the higher concentration (10 nM) we observed no P4-dependent enrichment (Fig. 4D and S4A). Therefore, the pulldown experiment replicates and supports the PLA2 activity data and shows that the interaction affinity of the two receptors is low in their apo state and increases significantly after they bind P4. Therefore, P4 shifts coreceptor affinity allowing their interaction at limiting concentrations to induce signaling.

### P4-mPRβ recruits ABHD2 from an intracellular compartment to the PM

We next wanted to translate the findings from the in vitro expression system to the oocyte. Defining the subcellular localization of ABHD2 was limited by the absence of good antibodies to detect endogenous ABHD2 and the fact that the Cherry-tagged ABHD2 constructs were not functional (Fig. S1). We therefore relied on surface biotinylation of PM proteins followed by streptavidin pulldown in oocytes expressing ABHD2-His (Fig. 5A). ABHD2-His is not detectable at the PM at rest, however it is enriched at the PM in response to P4 in a time dependent fashion (Fig. 5A). The amount of ABHD2 recovered by surface biotinylation increased between 5 and 30 min after P4 treatment (Fig. 5A).

**Figure 5.**
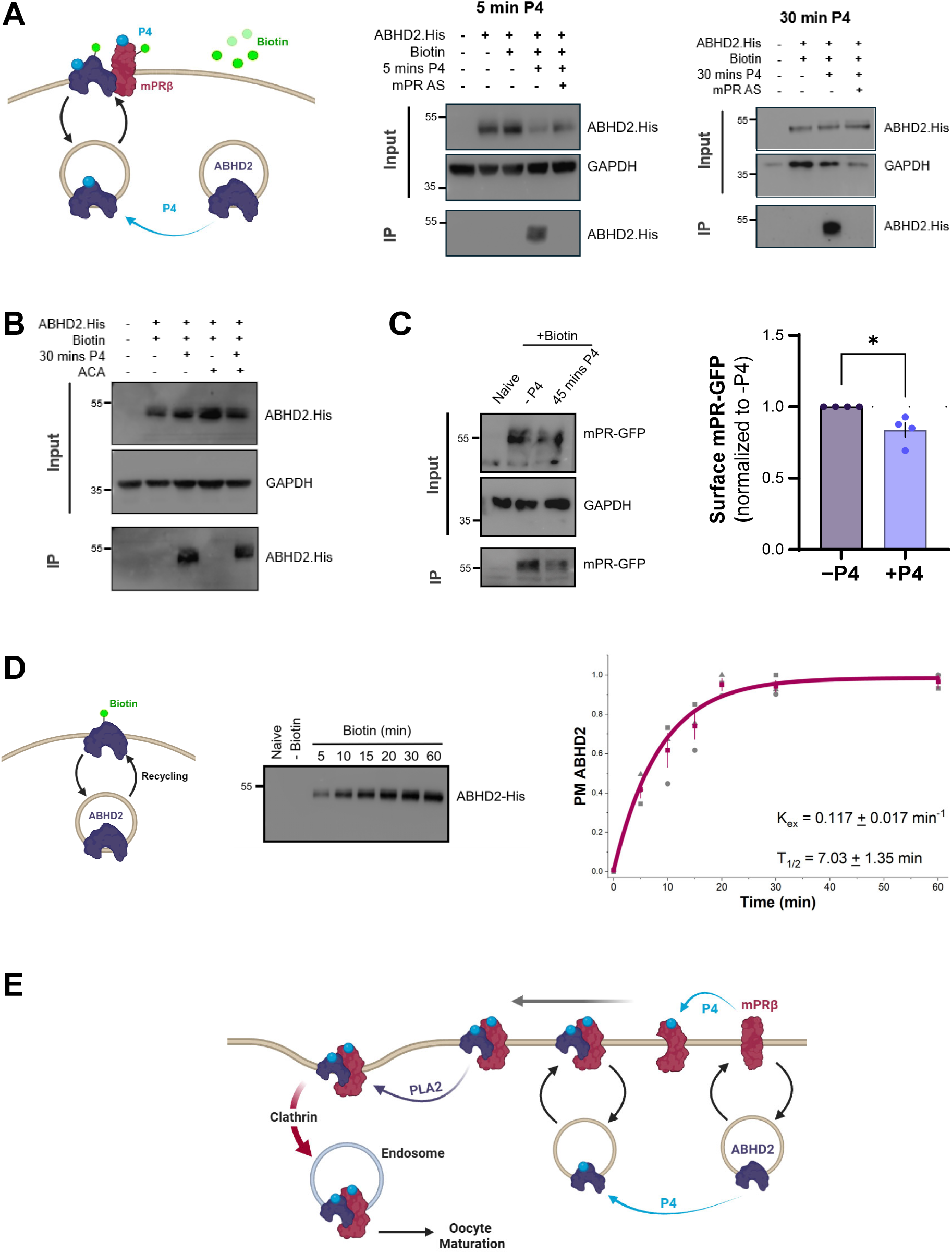
P4 promotes mPRβ-dependent PM accumulation of ABHD2. **(A)** Schematic representation and Western blots to quantify ABHD2 at the PM. Oocytes expressing ABHD2-His, with or without mPRβ antisense (mPR.AS), were treated with P4 (10^−5^ M) or ethanol for 5- or 30-min. Surface proteins were labeled with membrane-impermeant biotin, isolated by streptavidin pulldown, and analyzed by Western blotting. Representative input and streptavidin-bound fractions. Blots are representative of three to four independent experiments. **(B)** Oocytes expressing ABHD2-His were pretreated with the PLA, inhibitor N-(p-amylcinnamoyl) anthranilic acid (ACA; 50 µM) for 30 min before P4 stimulation (10^−5^ M) for an additional 30 min. Surface-biotinylated and pulldown were as in panel A. Blots are representative of three independent experiments. **(C)** Representative Western blots showing surface biotinylation of mPRβ-GFP in oocytes treated with P4 (10^−5^ M) or ethanol for 45 min. Input and streptavidin-bound fractions were analyzed by Western blotting, with GAPDH serving as a loading control. Blots are representative of four independent experiments. Surface mPRβ-GFP was quantified as the ratio of streptavidin-bound mPRβ-GFP following P4 treatment over before P4 treatment (+P4/−P4). Data are presented as Mean ± SEM, n = 4, unpaired t-test (* p < 0.05). **(D)** Schematic representation of ABHD2 biotinylation tagging. Oocytes expressing ABHD2-His were incubated with membrane-impermeant biotin (1 mg/mL) at room temperature for the indicated times (0–60 min), followed by streptavidin pulldown and Western blotting. ABHD2-His band intensities were quantified and normalized to the maximum intensity measured within each independent experiment. The time course of ABHD2 biotinylation was fitted using a one-phase exponential association model. The fit yielded an apparent exocytosis rate constant (K_ex_) of 0.117 ± 0.017 min^−1^ and a half-time (t½) of 7.03 ± 1.35 min. Data are presented as Mean ± SEM, n = 3. **(E)** Proposed model of rapid progesterone signaling in *Xenopus* oocytes.

Importantly the P4-dependent translocation of ABHD2 to the PM requires mPRβ as it is lost in oocytes where mPRβ is knocked down (Fig. 5A, mPR-AS). Furthermore, the translocation of ABHD2 to the PM in response to P4 is not dependent on its PLA2 activity as it was not blocked by ACA (Fig. 5B), a PLA2 inhibitor that blocks ABHD2 in the oocyte ^24^.

In contrast to ABHD2, mPR is detected at the PM at rest and its PM levels decrease 45 min after P4 addition (Fig. 5C), showing that mPR is internalized in response to P4, consistent with our previous findings ^24^.

These results argue that ABHD2 is enriched intracellularly at steady state but that it continuously cycles through the PM. P4 traps ABHD2 at the PM in an mPRβ-dependent fashion. To test this mechanistically, we assessed the surface accessibility of ABHD2 using the biotinylation assay in the absence of P4 treatment. If ABHD2 is constantly recycling through the PM, the intracellular ABHD2 population is expected to become surface accessible over time. Incubating oocytes with biotin extracellularly shows a time-dependent increase in the levels of biotinylated ABHD2 (Fig. 5D and S4B). The time course estimates an exocytosis rate of 0.117±0.017 min^-1^ and T_1/2_ of 7.03±1.35 min^-1^ (Fig. 5D and S4B). This confirms that although at steady state ABHD2 is intracellular, it is continuously sampling the PM. In response to P4 the affinity of mPRβ to ABHD2 increases, which allows mPRβ to trap the recycling ABHD2 population and inducing PLA2 activity, which leads to internalization of the coreceptor complex to release the oocyte meiotic arrest (Fig. 5E).

## Discussion

The oocyte’s commitment to maturation is one of the most critical processes in biology as it must be timed properly to produce a mature egg able to support fertilization and early embryonic development. P4 triggers oocyte maturation through a rapid signaling cascade as the fully grown oocyte is arrested in prophase of meiosis I and is transcriptionally silent. This mode of rapid P4 signaling has important roles in the reproductive, nervous, cardiovascular, immune, and metabolic system ^10–14^. Yet despite its increasingly recognized roles the P4-mPR signaling cascade is poorly defined.

There is strong evidence in the literature for a role for trimeric G-proteins in mPR signaling, especially for mPRα ^15^. It is however unclear whether this is direct with mPRs themselves functioning as GPCRs, or indirect with mPRs producing lipid messengers that then activate GPCRs ^24^. There is also much to be learned from the different receptor classes that mediate the P4 rapid action ^17^. The release of the oocyte meiotic arrest in the frog requires two classes of P4 receptors, mPRβ and ABHD2 ^24^. They work in coordination to induce a PLA2 activity that generates lipid second messengers, which in turn act on GPCRs within the oocyte. Blocking PLA2 activity, mutating the ABHD2 lipase catalytic domain, or knockdown of either mPRβ or ABHD2 all inhibit P4-induced oocyte maturation ^24^.

Here we define the early steps of interactions between mPRβ and ABHD2 to mediate the P4 signal. We show that at rest the two receptors localize to two different subcellular compartments: mPRβ is enriched at the PM, whereas ABHD2 is primarily intracellular. Despite their steady state distribution, both receptors recycle at the PM. This is evidenced for mPRβ by blocking exocytosis, which leads to its enrichment intracellularly and induction of maturation in a P4-independent manner ^23,29^. For ABHD2 surface biotin labeling of PM proteins at rest does not detect ABHD2 (Fig. 5A), whereas incubating oocytes with biotin for extended periods to allow for PM recycling labels increasing levels of ABHD2 in a time-dependent manner (Fig. 5D and S4B). We further show that P4 binds to both mPRβ and ABHD2 and increases their interaction affinity. This leads to trapping of ABHD2 at the PM through its interaction with mPRβ and the activation of the co-receptor PLA2 activity to release the oocyte meiotic arrest (Fig. 5E).

This represents a novel mode of signaling for PM receptors. Although several receptor classes rely on endocytosis during their signaling cycle ^31^, this is the only case we are aware of where two distinct receptors that bind to the same agonist employ a trafficking trap mechanism to activate the coreceptor complex. Both receptor tyrosine kinases and GPCRs are endocytosed following agonist binding, and endocytosis is even more critical for the Ser/Thr TGFβ receptors to allow them to interact with the SMAD proteins that reside in endosomes ^31^. In contrast, in the case of P4-mPRβ-ABHD2 signaling the two receptors are in distinct subcellular location and agonist binding enriches the complex at the PM through increased co-receptor interaction (Fig. 5E). Importantly the role of the agonist in this case is to allow the formation of the co-receptor complex at the PM by increasing their interaction affinity and not to induce the PLA2 activity, which is independent of P4 binding once the co-receptor complex is formed.

We further show that *Xenopus* ABHD2 is a two-pass transmembrane protein with intracellular N- and C-termini (Fig. 2). This allows access of the lipase catalytic domain to the core of the bilayer fatty acid tails. Although this topology fits well with ABHD2 function, it contradicts the known structure of other members of the ABHD family and the predicted Alphafold2 structure of mammalian ABHD2, which is primarily formed of β-sheets, including in the area of the second TM domain. We tested ABHD2 topology in cells and in a microsome expression system, both of which support a two-TM topology. It is possible that unlike other ABHD members ABHD2 adopts a different confirmation within the bilayer. Resolving these discrepancies would require an atomic solution structure of ABHD2.

Our data with the biotin tagged P4 analogues argue that *Xenopus* ABHD2 binds P4. This conclusion is supported by functional data that show that the P4-Biotin with the long linker binds microsome expressed ABHD2 and induces oocyte maturation, whereas the one with the short linker does not bind ABHD2 nor does it induce maturation. These findings are consistent with those of Miller et al. who also concluded that P4 binds ABHD2 ^18^. Recently Edwards et al. tested the ability of human ABHD2 to bind P4 and other steroids ^27^. They used a recombinant full length or truncated ABHD2 missing the first TM domain purified from Sf9 cells in a differential scanning fluorimetry assay and observed no binding with P4. The reasons for these discrepancies are unclear but could be due to the purification methods or the binding assays employed.

Collectively, our results elucidate a novel mechanism of rapid steroid signaling. P4 signaling in the oocyte requires both mPRβ and ABHD2. ABHD2 is enriched intracellularly at steady state whereas mPRβ is enriched at the PM (Fig. 5E). P4 binds both receptors and increases their interaction affinity to stabilize the coreceptor complex at the PM and activate PLA2 (Fig. 5E). This then supports coreceptor internalization and the release of the oocyte meiotic arrest.

## Materials and Methods

### Reagents

Antibodies, chemicals, and other reagents are listed in Supplementary Table 1.

### Xenopus laevis oocytes

All animal procedures were approved by the Weill Cornell Medicine–Qatar (WCM-Q) Institutional Animal Care and Use Committee. Adult female Xenopus laevis frogs were obtained from Xenopus 1, and stage VI oocytes were isolated as previously described ³². Oocytes were maintained at 18°C in L-15 medium (Sigma-Aldrich, Cat. No. L4386) supplemented with HEPES (Sigma-Aldrich, Cat. No. H4034), 0.1% penicillin–streptomycin (Thermo Fisher Scientific, Cat. No. 15140-122), and 0.1% gentamicin (EMD Millipore, Cat. No. 345814-1GM), pH 7.6. Oocytes were used 24–72 h after isolation. Germinal vesicle breakdown (GVBD) was assessed by the appearance of a white spot at the animal pole.

### Cell culture and transfection

HeLa cells were maintained in Dulbecco’s modified Eagle’s medium (DMEM) supplemented with 10% heat-inactivated fetal bovine serum (FBS; Thermo Fisher Scientific, Cat. No. 10082147) and 1% penicillin–streptomycin at 37°C in a humidified atmosphere containing 5% CO,. Cells were co-transfected with plasmids encoding Orai1-mCherry (0.7 µg) and N- or C-terminally His-tagged ABHD2 (0.5 µg) using Lipofectamine 2000 (Thermo Fisher Scientific, Cat. No. 11668030) in Opti-MEM. After 7 h, the transfection medium was replaced with complete DMEM. Cells were fixed 24 hours after transfection.

### Molecular biology

The pSGEM-mPRβ-GFP construct was described previously²³. The *Xenopus laevis* ABHD2.S coding sequence (Xenbase ID: XB-GENE-17345096) was commercially synthesized by Gene Universal. Constructs encoding ABHD2 with an N-terminal (His-ABHD2) or C-terminal (ABHD2-His) 6×His tag were generated in the pSGEM expression vector and pcDNA3.1(−) by Gene Universal. Both His-tagged ABHD2 constructs contained EcoRI and HindIII restriction sites flanking their coding sequences.

To generate C-terminally His-tagged mPRβ (mPRβ-His) in pALiCE02, mPRβ was amplified using the following primers:

Forward: 5′-CTACCATGGCAATGACTACCGCAATCCTTGA-3′

Reverse: 5′-CTAGGTACCTCAGTGATGGTGATGGTGATGAAGTTCTTTTCTGGCCAACT-3′

The PCR product was digested with NcoI and KpnI and ligated into pALiCE02 digested with the same restriction enzymes to remove the in-frame YFP tag.

To generate C-terminally His-tagged ABHD2 (ABHD2-His) in pALiCE02, ABHD2 was amplified using the following primers:

Forward: 5′-ACTGATATCATGGATGCGATAGTGGAAACCC-3′

Reverse: 5′-CTAGGTACCTCAGTGATGGTGATGGTGATGTTTATGGTCAGACTCAGCAGCC-3′

The PCR product was digested with EcoRV and KpnI and ligated into pALiCE02 that had been digested with NcoI, blunt-ended using the Klenow fragment to generate an EcoRV-compatible end and subsequently digested with KpnI to remove the in-frame YFP tag.

ABHD2 construct (His-ABHD2) and a C-terminally HA-tagged mPRβ construct (mPRβ-HA) were commercially synthesized and cloned into pALICE02 by Gene Universal. All constructs were verified by sequencing and restriction digestion. For *Xenopus laevis* oocyte expression, pSGEM constructs were linearized with NheI and transcribed in vitro using the mMESSAGE mMACHINE T7 Transcription Kit (Thermo Fisher Scientific, Cat. No. AM1344).

### ALiCE cell-free protein expression

Proteins were expressed using the ALiCE Maxi Kit (LenioBio GmbH, Cat. No. AL0103500). pALiCE02 plasmids were added at the indicated final DNA concentrations, and reactions were incubated for 48 h with shaking at 700 rpm at 25°C. Protein expression was confirmed by Western blotting.

### Immunoprecipitation

For ALiCE samples, reactions were diluted 1:200 in immunoprecipitation buffer containing 30 mM HEPES and 100 mM NaCl (pH 7.5), and 20 µL was retained as input. For HA immunoprecipitation, ALiCE samples co-expressing ABHD2 and mPRβ were divided into two aliquots and treated with either unlabeled progesterone (P4; 10^−5^ M) or ethanol vehicle for 30 min at 4°C to prevent vesicular trafficking. Following treatment, Nonidet P-40 Substitute (Sigma-Aldrich, Cat. No. 74385) was added to both conditions at a final concentration of 0.5% to disrupt membrane vesicles, and samples were mixed by pipetting. HA-tagged proteins were immunoprecipitated using 50 µL Pierce Anti-HA Magnetic Beads (Thermo Fisher Scientific, Cat. No. 88837) and eluted in 50 µL 2× Laemmli buffer.

For His immunoprecipitation, samples were maintained in detergent-free buffer to preserve membrane vesicle integrity. His-tagged proteins were immunoprecipitated using 50 µL µMACS Anti-His MicroBeads (Miltenyi Biotec, Cat. No. 130-091-124) and µ Columns, followed by elution in 50 µL preheated elution buffer supplied with the kit.

### Cell-surface Biotinylation

Oocytes expressing ABHD2-His for 48 h, with or without mPR antisense injection, were treated with 10^−5^ M P4 or ethanol for 5 or 30 min. For 30-min treatments, oocytes were maintained at room temperature for 25 min and transferred to 4°C for the final 5 min. Surface proteins were labeled with 1 mg/mL Sulfo-NHS-SS-Biotin (Thermo Fisher Scientific, Cat. No. 21331) for 30 min at 4°C, quenched with 50 mM Tris-HCl, and washed with Ringer’s solution. Oocytes were lysed as described above, and biotinylated proteins were isolated using Pierce Streptavidin Magnetic Beads (Thermo Fisher Scientific, Cat. No. 88817) for 1 h at 4°C. Bound proteins were eluted in 50 µL 2× Laemmli buffer and analyzed by Western blotting.

For the time course, oocytes were incubated with Sulfo-NHS-SS-Biotin for 5, 10, 15, 20, 30, or 60 min at room temperature. To estimate the K_ex_ and the T_1/2_ of ABHD2 recycling the data was fitted in Origin 2026 using to a one phase exponential growth function:

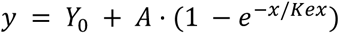

Where Y_0_ represents the basal PM ABHD2 levels at time 0; A the maximal biotin labelled ABHD2 when the entire ABHD2 cellular pool is labelled; and K_ex_ (min^-1^) the exocytosis rate constant, which reports the proportion of the intracellular ABHD2 pool delivered to the PM per minute. The fit yielded an R^2^ = 0.9976.

To inhibit PLA, activity, oocytes were pretreated with 50 µM N-(p-amylcinnamoyl) anthranilic acid (ACA; Sigma-Aldrich, Cat. No. A8486) for 30 min. P4 (10^−5^ M) was then added for an additional 30 min without removing ACA. During P4 treatment, oocytes were maintained at room temperature for 25 min and transferred to 4°C for the final 5 min before surface biotinylation.

To assess P4-induced changes in surface mPRβ, oocytes expressing mPRβ-GFP were treated with P4 (10^−5^ M) or ethanol vehicle for 45 min and transferred to 4°C before surface biotinylation. Biotinylated proteins were isolated using streptavidin magnetic beads and analyzed by Western blotting as described above.

### Progesterone-biotin binding assay

ABHD2 and mPRβ were expressed separately using the ALiCE cell-free protein expression system. Expression reactions were diluted 1:200 in binding buffer containing 30 mM HEPES and 100 mM NaCl (pH 7.5), and 20 µL was retained as input. The remaining samples were divided into 490-µL aliquots and incubated with progesterone-3-biotin (P4-3-B; Cayman Chemical, Cat. No. 9000644), progesterone-3-PEG11-biotin (P4-PEG-B; Cayman Chemical, Cat. No. 9000645), or ethanol vehicle for 30 min. Both biotinylated progesterone analogues were used at a final concentration of 10^−5^ M.

Pierce Streptavidin Magnetic Beads (Thermo Fisher Scientific, Cat. No. 88817) were blocked with 3% BSA and 0.2% Tween-20 for 30 min, washed with binding buffer, and incubated with the samples for 30 min. Beads were then washed sequentially for 15 min with: (i) 1 M NaCl, 1% Ecosurf EH-9, 0.5% sodium deoxycholate, 0.1% SDS, and 50 mM Tris-HCl (pH 8.0); (ii) 0.5 M NaCl, 30 mM HEPES, and 0.2% Triton X-100; and (iii) 20 mM Tris-HCl (pH 7.5). Bound proteins were eluted in 50 µL 2× Laemmli buffer and analyzed by Western blotting.

For competitive elution experiments, following streptavidin pulldown and washing, beads were incubated with free progesterone (P4; 10^−5^ or 10^−4^ M) for 1h at RT. The supernatants containing proteins released during P4 incubation were collected and mixed with 2× Laemmli buffer. Proteins remaining associated with the beads were subsequently eluted using 2× Laemmli buffer. Both fractions were analyzed by Western blotting.

### Western blotting

Oocytes were lysed in MPF buffer containing 80 mM β-glycerophosphate, 20 mM HEPES (pH 7.5), 15 mM MgCl,, 20 mM EGTA, 1 mM sodium orthovanadate, 50 mM NaF, 1 mM DTT, and protease and phosphatase inhibitors. Yolk was removed by three consecutive centrifugations at 1,000 × g for 10 min at 4°C. Proteins were separated by 10% SDS-PAGE and transferred onto PVDF membranes. Membranes were blocked for 1 h in 5% nonfat milk prepared in Tris-buffered saline containing Tween-20 (TBS-T), followed by overnight incubation at 4°C with primary antibodies diluted in 3% BSA/TBS-T. HRP-conjugated or IRDye 680/800-conjugated secondary antibodies were used at a dilution of 1:5,000. Chemiluminescent signals were detected using ECL and quantified using ImageJ. Fluorescent signals were acquired using a LI-COR Odyssey CLx imaging system and analyzed using Image Studio software.

Primary antibodies were used at the following dilutions: anti-ABHD2, anti-GFP, anti-phospho-Cdc2, anti-His, and anti-mPRβ, 1:1,000; anti-GAPDH and anti-tubulin, 1:10,000; and anti-phospho-MAPK, 1:4,000.

### Phospholipase A, assays

The indicated final DNA concentrations of mPRβ and/or ABHD2 constructs were added to the ALiCE cell-free protein expression system. Where indicated, microsomes containing separately expressed proteins were mixed and incubated with 10% polyethylene glycol (PEG; Sigma-Aldrich, Cat. No. P7181) for 15 min at 37°C to promote membrane fusion. Expression of mPRβ and ABHD2 was confirmed by Western blotting. ALiCE reactions without added DNA served as background controls. Samples were treated with P4 (10^−5^ M) or ethanol.

PLA, activity was assessed using the Phospholipase A, Activity Assay Kit (Fluorometric; Abcam, Cat. No. ab273278). Fluorescence was recorded for 60 min using a CLARIOstar plate reader (BMG LABTECH) at excitation and emission wavelengths of 388 and 513 nm, respectively, at 37°C. Fluorescence measurements were converted to nmol of product using a standard curve generated with the fluorescence standard supplied in the kit. Background values from ALiCE reactions without added DNA were subtracted from the corresponding experimental measurements. Technical replicates (3–4 wells per condition) were averaged within each independent experiment.

For live-oocyte imaging, the fluorescent PLA, substrate PED6 (Thermo Fisher Scientific, Cat. No. D23739) was microinjected into *Xenopus laevis* oocytes to achieve an estimated intracellular concentration of 10 µM. Oocytes were immobilized using cyanoacrylate adhesive, with the animal pole facing the objective, and imaged at 5-min intervals using an upright Zeiss confocal microscope equipped with a 20× water-immersion objective. PED6 fluorescence was excited using the 488-nm laser line, and emission was collected between 493 and 598 nm. Following baseline acquisition, P4 (10^−5^ M) or ethanol vehicle was added. Mean fluorescence intensity was quantified using Fiji/ImageJ and normalized to the pretreatment baseline of each oocyte.

### Immunofluorescence

HeLa cells were fixed with 1% paraformaldehyde (PFA). For membrane permeabilization, cells were treated with 0.3% Triton X-100. For selective plasma membrane permeabilization, cells were treated with 0.01% digitonin in NHM buffer (20 mM HEPES, 110 mM NaCl, and 2 mM MgCl,, pH 7.4) for 5 min. Nonpermeabilized control cells were processed without detergent treatment. Cells were incubated overnight with rabbit anti-His antibody (LSBio, Cat. No. LS-C60077-100) or rabbit anti-STIM1 antibody (Cell Signaling Technology, Cat. No. 5668), each diluted 1:500, followed by incubation with Alexa Fluor 488-conjugated anti-rabbit secondary antibody for 2 h. Nuclei were stained with Hoechst 33342 (Thermo Fisher Scientific, Cat. No. H3570; 1:5,000) for 5 min. Images were acquired using a Zeiss LSM 880 confocal microscope equipped with a Plan-Apochromat 63×/1.4 NA oil-immersion objective and analyzed using Fiji/ImageJ.

### Statistics

Data are presented as mean ± SEM from at least three independent experiments. Statistical analyses were performed using GraphPad Prism 9, with the tests indicated in the figure legends. Statistical significance is indicated as follows: ns, not significant; * *p* ≤ 0.05; ** *p* ≤ 0.01; and *** *p* ≤ 0.001.

## Materials availability statement

All clones or reagents generated in the course of this study are available to colleagues in the scientific community without restrictions. If interested please contact the corresponding author, Khaled Machaca at.

## Acknowledgments

We are grateful to the Vivarium and Microscopy Cores at WCMQ for their support in multiple experiments. This work as well as the Cores are supported by the Biomedical Research Program at Weill Cornell Medical College in Qatar (BMRP), a program funded by Qatar Foundation. The findings herein reflect the work and are solely the responsibility of the authors.

## Supplementary Materials

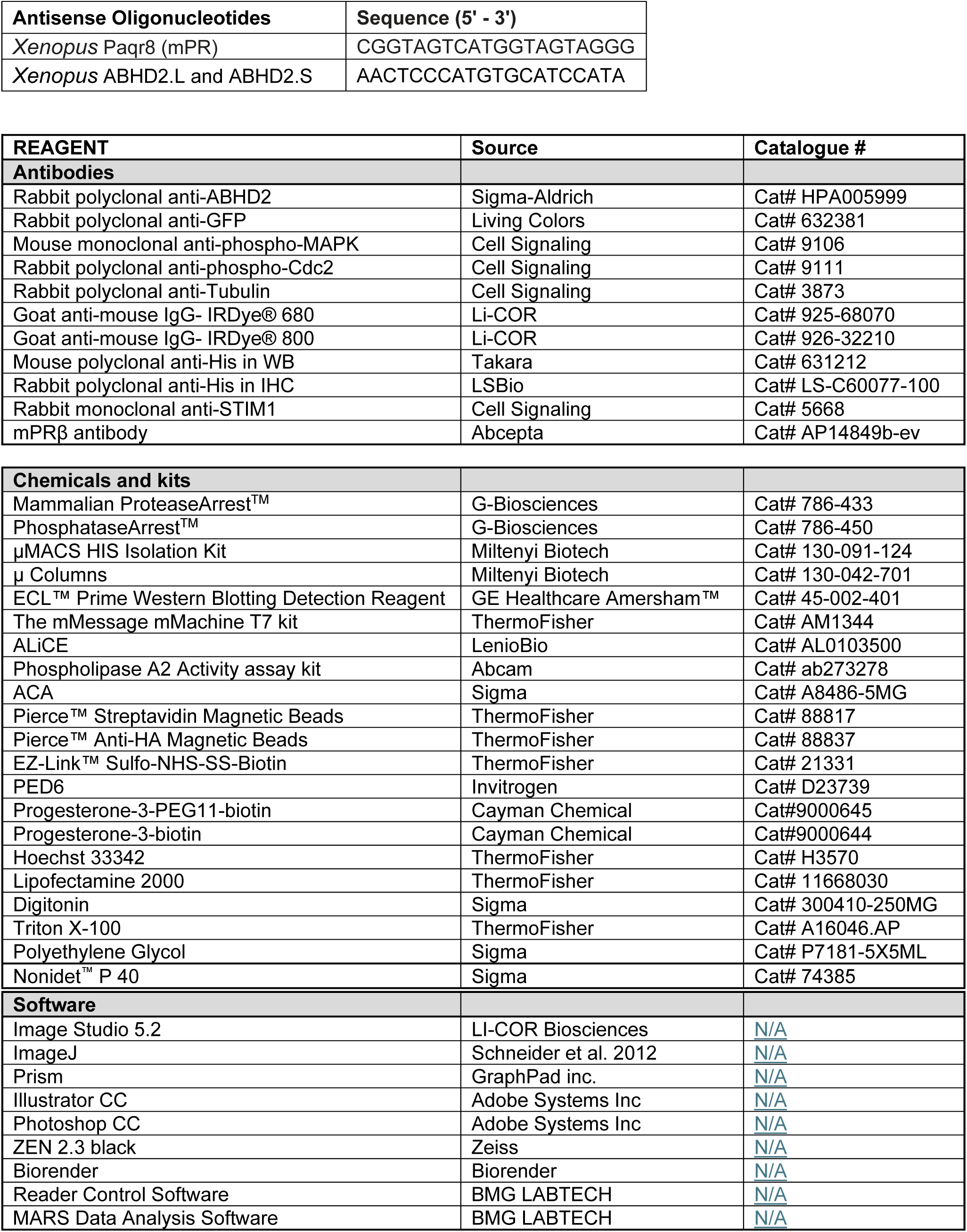

## Supplementary Figure Legends

**Supplementary Figure 1.**
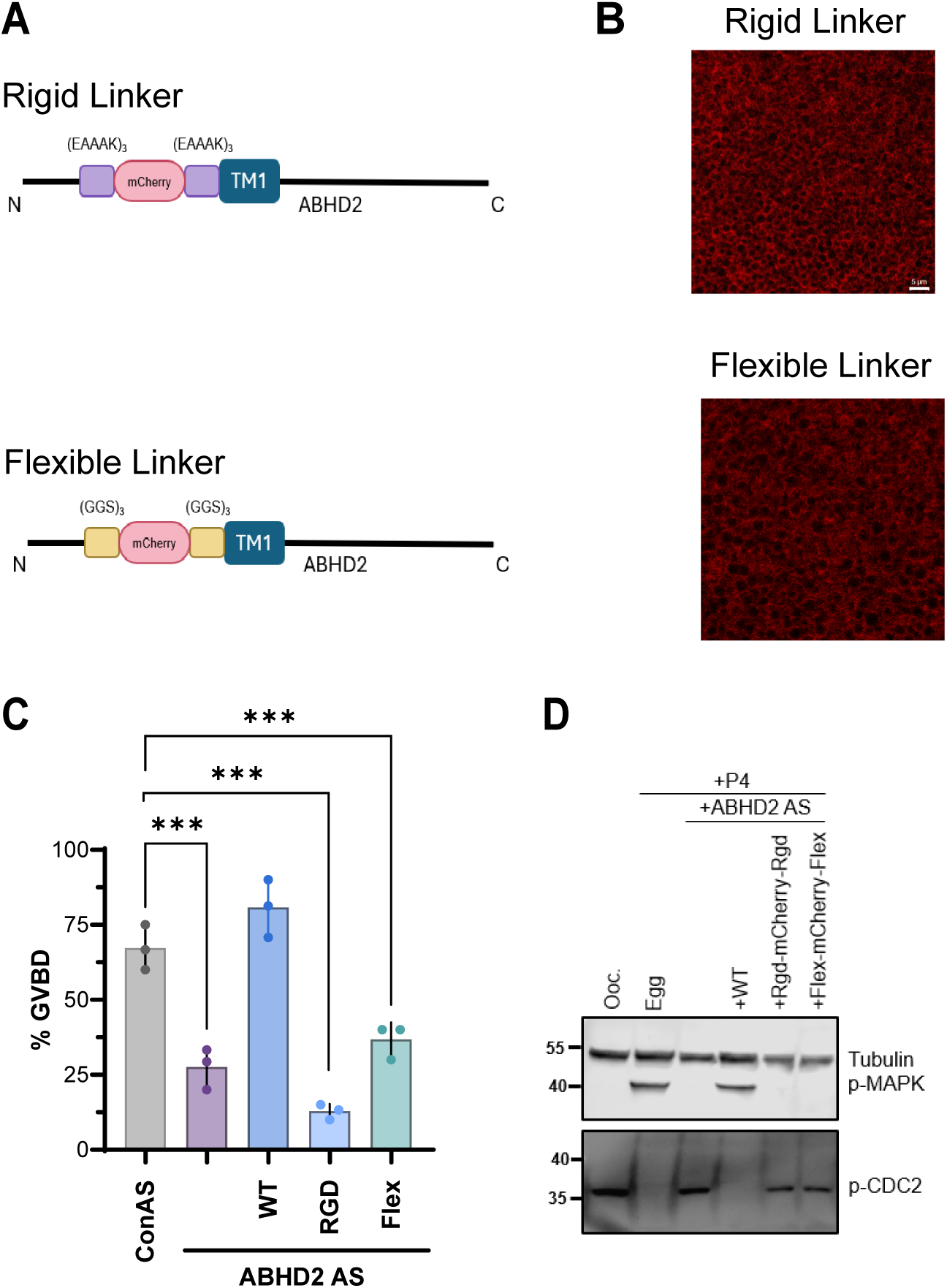
ABHD2 does not tolerate an mCherry-tag. **(A)** Schematic representation of *Xenopus* ABHD2 constructs containing mCherry inserted immediately upstream of the first transmembrane domain (TM1), flanked by either rigid (EAAAK)_3_ or flexible (GGS)_3_ linkers. The constructs are designated ABHD2-Rigid-mCherry-Rigid and ABHD2-Flex-mCherry-Flex, respectively. **(B)** Representative confocal fluorescence images of oocytes expressing ABHD2-Rigid-mCherry-Rigid or ABHD2-Flex-mCherry-Flex. Images represent maximum-intensity projections of Z-stacks generated using ZEN software. Red fluorescence indicates expression of the mCherry-tagged constructs. Scale bar 5 µm. **(C)** Percent germinal vesicle breakdown (GVBD) in control antisense (Con AS) or ABHD2 antisense (ABHD2 AS) injected oocytes expressing wild-type ABHD2 (WT), ABHD2-Rigid-mCherry-Rigid, or ABHD2-Flex-mCherry-Flex following P4 stimulation. Data are presented as Mean ± SEM, n = 3, one-way ANOVA with Dunnett correction (*** p < 0.001). **(D)** Representative Western blots showing P4-induced MAPK phosphorylation and Cdc2 dephosphorylation in control oocytes and ABHD2 antisense (ABHD2 AS)-injected oocytes expressing wild-type ABHD2 or the mCherry-tagged constructs. Tubulin serves as loading control. Representative of three independent experiments.

**Supplementary Figure 2.**
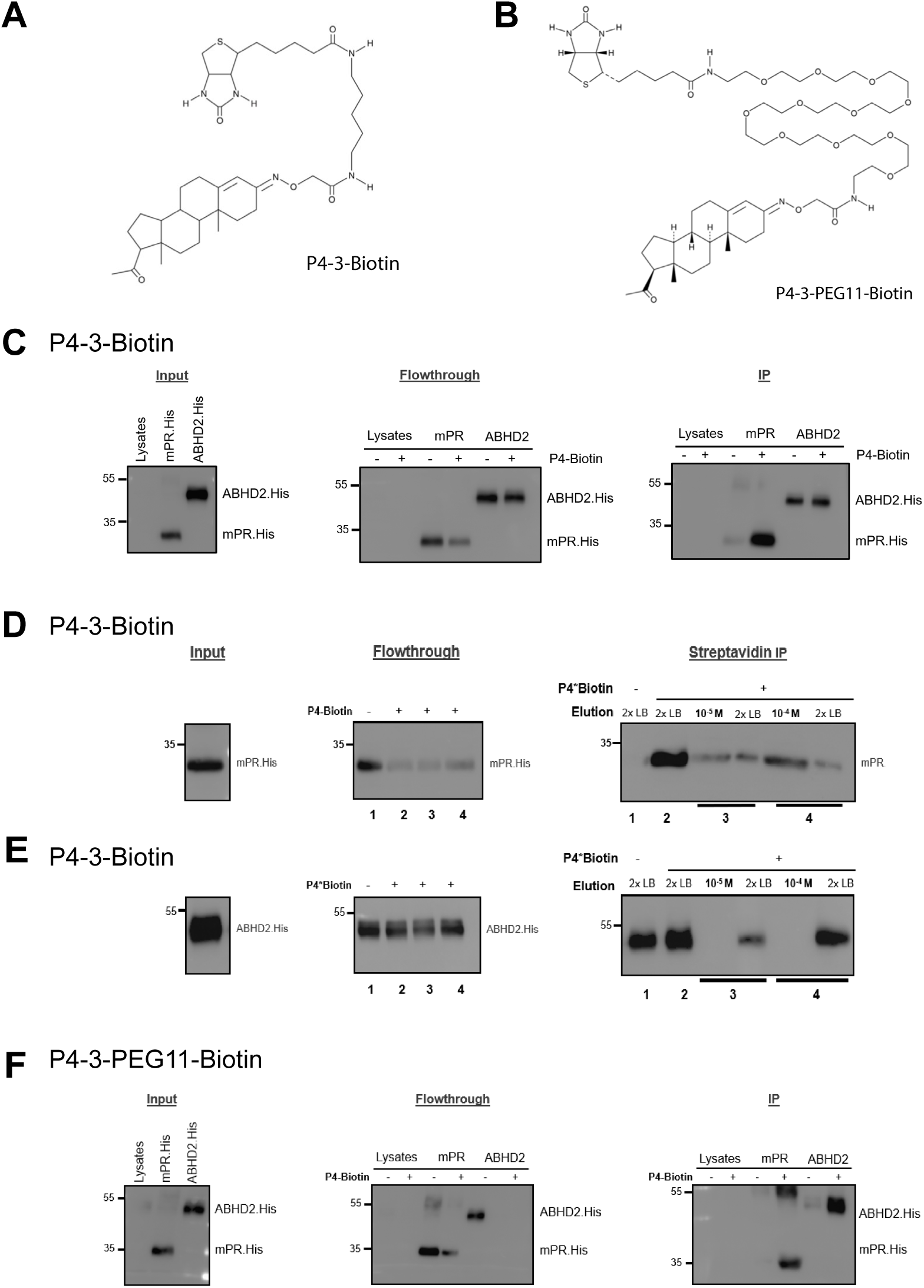
Binding of biotinylated P4 analogues to mPRβ and ABHD2. **(A)** Chemical structure of P4-3-Biotin containing a short linker between P4 and biotin. **(B)** Chemical structure of P4-3-PEG11-Biotin containing a long linker between P4 biotin. **(C)** Binding of P4-3-Biotin to mPRβ or ABHD2 expressed in microsomes. Microsomes expressing His-tagged mPRβ or ABHD2 were incubated with P4-3-Biotin (10^−5^ M) or ethanol, followed by streptavidin pulldown and Western blotting using an anti-His antibody. Representative Western blots of input, flow-through, and streptavidin-bound fractions are shown. Blots are representative of three independent experiments. **(D-E)** Competitive elution assay examining the specificity of P4-3-Biotin binding to mPRβ (D) and ABHD2 (E). Following incubation of microsomes expressing the different proteins with P4-3-Biotin and capture on streptavidin beads, samples were washed and proteins eluted with free P4 (10^−5^ or 10^−4^ M). The P4-containing supernatants were collected and proteins remaining associated with the beads were subsequently eluted using 2× Laemmli buffer (2x LB). Representative Western blots of input, flow-through, and fractions recovered following the indicated elution conditions are shown. Blots are representative of three independent experiments. **(F)** Binding of P4-3-PEG11-Biotin to mPRβ or ABHD2 expressed in microsomes. Microsomes were incubated with P4-3-PEG11-Biotin (10^−5^ M) or ethanol, followed by streptavidin pulldown and Western blotting using an anti-His antibody. Representative Western blots of input, flow-through, and streptavidin-bound fractions are shown. Blots are representative of three independent experiments.

**Supplementary Figure 3.**
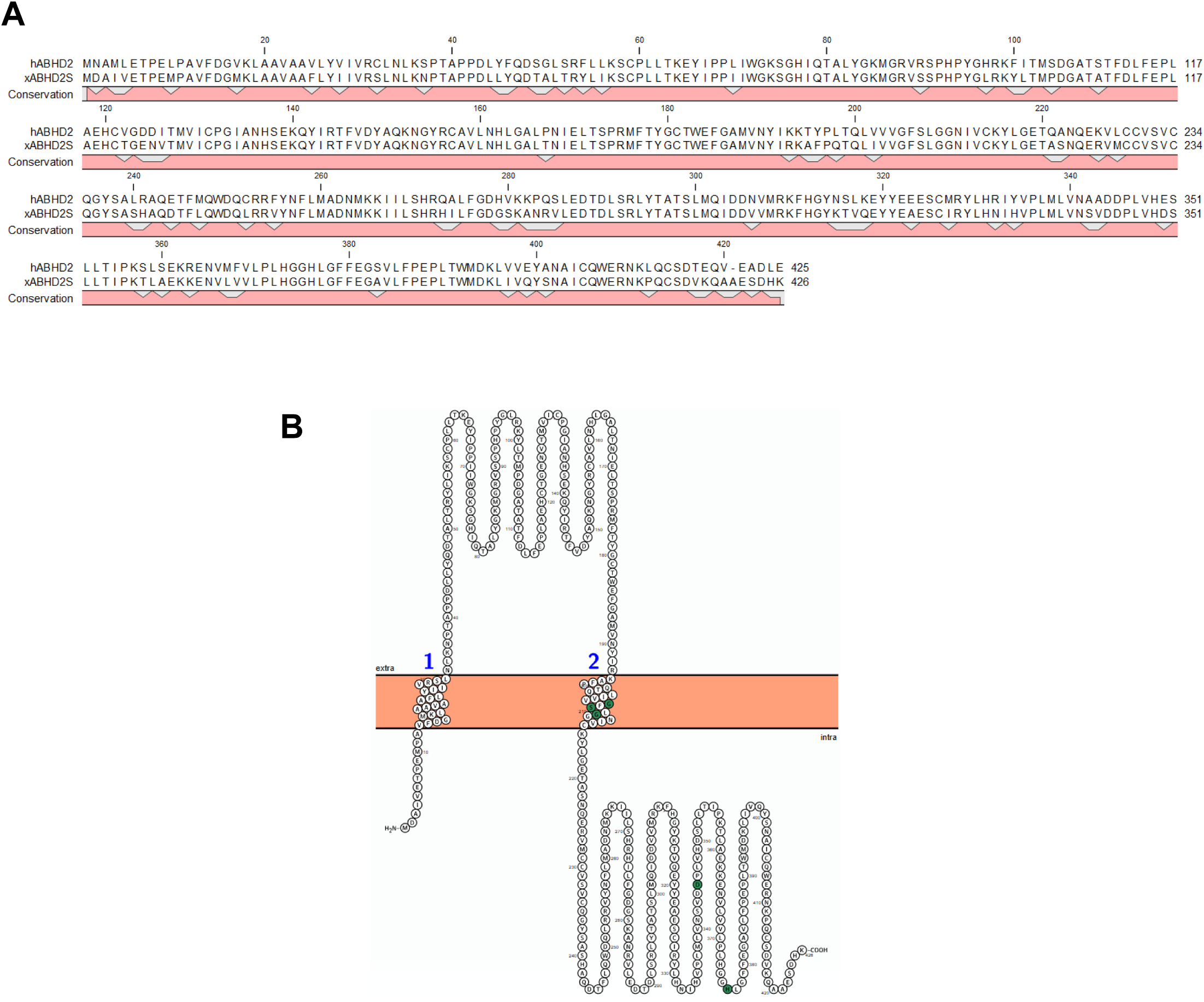
Sequence alignment and membrane topology of ABHD2. **(A)** Amino acid sequence alignment of human ABHD2 (hABHD2) and *Xenopus laevis* ABHD2 (xABHD2), highlighting the conservation of amino acid residues between the two orthologs. Conserved residues are indicated by the conservation track beneath the aligned sequences. **(B)** Predicted membrane topology of ABHD2 illustrating the arrangement of putative transmembrane regions within the lipid bilayer.

**Supplementary Figure 4.**
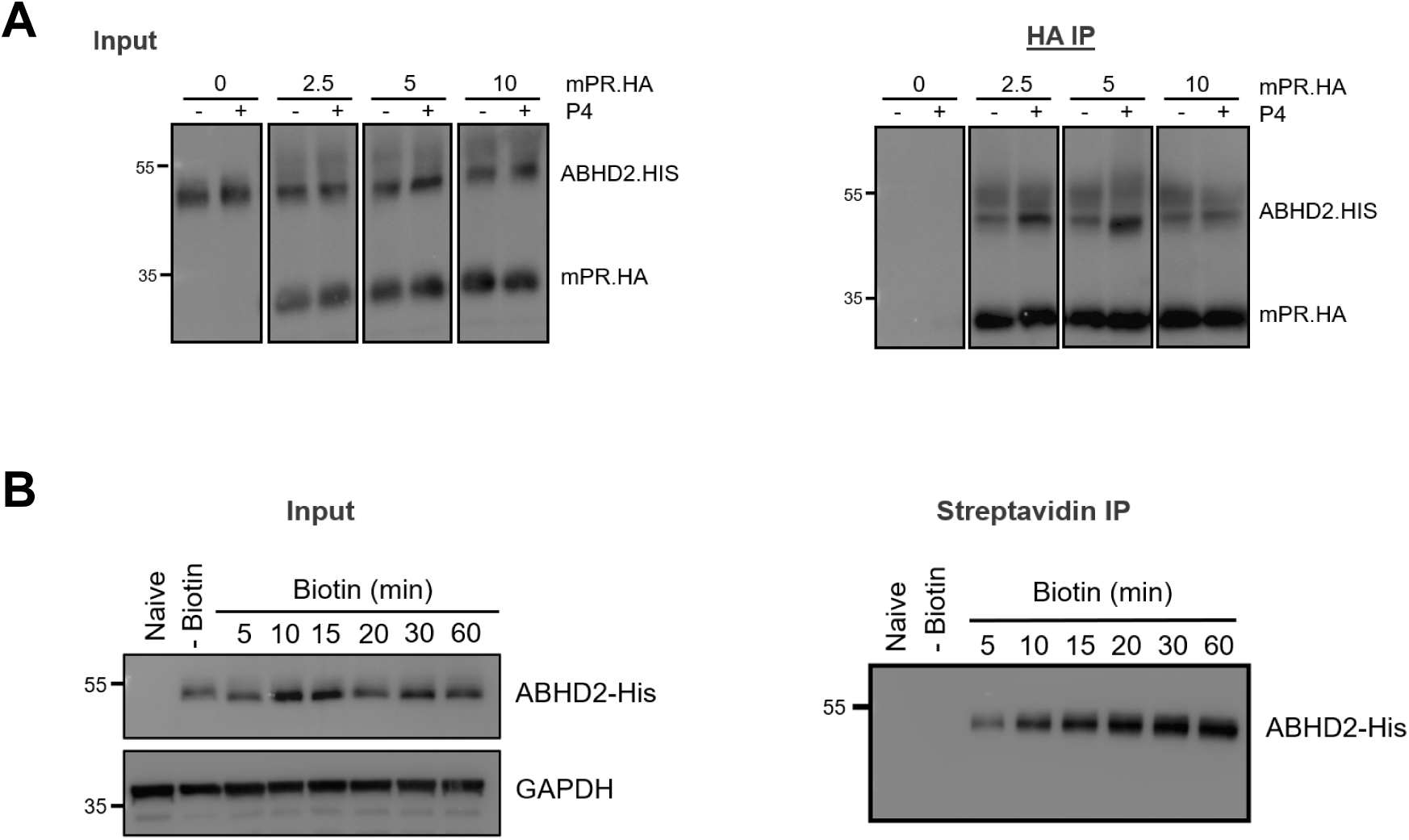
mPRβ–ABHD2 co-IP and ABHD2 PM recycling. **(A)** Representative Western blots of input fractions and co-immunoprecipitation for the experiment shown in Figure 4D. ABHD2-His was co-expressed with HA-tagged mPRβ at increasing mPRβ DNA concentrations (0, 2.5, 5, and 10 nM), while maintaining a constant ABHD2 DNA concentration (2.5 nM), in the ALiCE cell-free expression system. Microsomes were treated with progesterone (P4; 10^−5^ M) or ethanol. Blots are representative of five independent experiments. **(B)** Representative Western blots of input and streptavidin IP over the ABHD2 surface-biotinylation time course shown in Figure 5D. Oocytes expressing ABHD2-His were incubated with membrane-impermeant biotin (1 mg/mL) at room temperature for the indicated times (5–60 min), followed Western blotting. Naïve and non-biotinylated oocytes served as controls. GAPDH was used as a loading control. Blots are representative of three independent experiments using oocytes from three different frogs.

